# Incomplete dosage balance and dosage compensation in the ZZ/ZW Gila monster (*Heloderma suspectum*) revealed by *de novo* genome assembly

**DOI:** 10.1101/2023.04.26.538436

**Authors:** Timothy H. Webster, Annika Vannan, Brendan J. Pinto, Grant Denbrock, Matheo Morales, Greer A. Dolby, Ian T. Fiddes, Dale F. DeNardo, Melissa A. Wilson

**Affiliations:** Department of Anthropology, University of Utah, Salt Lake City, UT; School of Life Sciences, Arizona State University, Tempe, AZ; Center for Evolution and Medicine, Arizona State University, Tempe, AZ; Department of Zoology, Milwaukee Public Museum, Milwaukee, WI USA; Department of Genetics, Yale University, New Haven, CT; Center for Mechanisms of Evolution, Biodesign Institute, Tempe, AZ; 10x Genomics, Pleasanton, CA

**Keywords:** sex chromosome evolution, ZW system, squamate, Anguimorpha, genome assembly, expression, reptiles

## Abstract

Reptiles exhibit a variety of modes of sex determination, including both temperature-dependent and genetic mechanisms. Among those species with genetic sex determination, sex chromosomes of varying heterogamety (XX/XY and ZZ/ZW) have been observed with different degrees of differentiation. Karyotype studies have demonstrated that Gila monsters (*Heloderma suspectum*) have ZZ/ZW sex determination and this system is likely homologous to the ZZ/ZW system in the Komodo dragon (*Varanus komodoensis*), but little else is known about their sex chromosomes. Here, we report the assembly and analysis of the Gila monster genome. We generated a *de novo* draft genome assembly for a male using 10X Genomics technology. We further generated and analyzed short-read whole genome sequencing and whole transcriptome sequencing data for three males and three females. By comparing female and male genomic data, we identified four putative Z-chromosome scaffolds. These putative Z-chromosome scaffolds are homologous to Z-linked scaffolds identified in the Komodo dragon. Further, by analyzing RNAseq data, we observed evidence of incomplete dosage compensation between the Gila monster Z chromosome and autosomes and a lack of balance in Z-linked expression between the sexes. In particular, we observe lower expression of the Z in females (ZW) than males (ZZ) on a global basis, though we find evidence suggesting local gene-by-gene compensation. This pattern has been observed in most other ZZ/ZW systems studied to date and may represent a general pattern for female heterogamety in vertebrates.

## Introduction

The 11,302 recognized extant species of squamate reptiles, lizards and snakes, (Uetz et al., 2021) exhibit remarkable diversity in morphology, ecology, life history, physiology, and behavior (Sites et al., 2011). In particular, modes of sex determination abound in squamates and include temperature-dependent sex determination, male heterogamety (XX/XY sex determination, in which males have an X and a Y and females have two X chromosomes), and female heterogamety (ZZ/ZW genetic sex determination in which females have a Z and a W and males have two Z chromosomes), as well as a combination of multiple modes (Cornejo-Páramo et al., 2020; Gamble et al., 2015; Hill et al., 2018; Pennell et al., 2018; Pokorna & Kratochvíl, 2009; Quinn et al., 2007; Shine et al., 2002). The incredible number of transitions in sex determination combined with mosaic patterns of both rapid turnover and relative stasis in the squamate tree make this group ideally suited for understanding aspects of sex chromosome evolution (Gamble et al., 2015, 2017).

Despite this extraordinary diversity, some groups are characterized by stability (e.g., (Augstenová, Pensabene, Veselý, et al., 2021). The suborder Anguimorpha contains at least 239 species across seven families (Uetz et al., 2021), yet a series of recent studies suggest that the dominant mode of sex determination in this clade is a ZZ/ZW genetic system, and that sex chromosomes across this clade are likely homologous (Augstenová, Pensabene, Kratochvíl, et al., 2021; Iannucci et al., 2019; Johnson Pokorná et al., 2014; Rovatsos et al., 2019). If true, these sex chromosomes are among the oldest and most stable in amniotes (Rovatsos et al., 2019). Dating back to at least 115-180 mya (Zheng & Wiens, 2016), this system is comparable in age to therian mammals (Graves, 2016b; Wilson & Makova, 2009).

One important consequence of the evolution of differentiated sex chromosomes from an ancestral autosomal pair is a difference in gene copy number between males and females (e.g., ZZ versus ZW), leading to an imbalance in gene expression between the sexes. Because deviations from gene dosage balance can have profound and often deleterious phenotypic effects (Birchler et al., 2005); “dosage balance” referring to equal expression between the sexes, and, “dosage compensation” is expected to evolve via stabilizing selection to maintain expression levels of the Z (or X) in the heterogametic sex relative to the ancestral autosomal pair, may then also evolve (Gu & Walters, 2017). Despite these expectations, dosage balance (between male and female Z-linked transcripts) and dosage compensation (between the Z and ancestral autosomal state) are not universal and substantially vary among taxa in both completeness and mechanism (Graves, 2016a; Gu & Walters, 2017; Vicoso & Bachtrog, 2009). Perhaps most striking is the difference between male and female heterogametic systems: dosage balance and/or compensation are often observed in male heterogametic systems (XX/XY), while nearly all female heterogametic (ZZ/ZW) systems studied to date—other than Lepidoptera and a species of brine shrimp (Gu et al., 2019; Huylmans et al., 2017, 2019; Walters et al., 2015)—exhibit a lack of dosage balance and incomplete dosage compensation (Gu & Walters, 2017). While putative mechanisms have been proposed to explain this difference (Mullon et al., 2015), dosage compensation has been studied in very few squamates and work in additional taxa is needed to better understand its evolution (Pinto et al., 2023).

In this study, funded in part through a successful crowdfunding campaign (Wilson, 2019), we sequenced a high-quality *de novo* genome for the Gila monster (*Heloderma suspectum*; Figure 1) and generated additional genomic and transcriptomic data from three males and three females to better understand squamate, specifically anguimorph, sex chromosome evolution. Previous studies in anguimorph sex chromosome evolution have identified chicken (*Gallus gallus*) chromosome 28 as the homologous linkage group to the sex chromosome system in varanids, *Abronia*, and helodermatids (Rovatsos et al., 2019). Thus, we (1) investigated whether the ZZ/ZW chromosomes observed across Anguimorpha show evidence of homology at the genomic sequence level, representing a single, ancient evolutionary origin with subsequent losses in some lineages (Rovatsos et al., 2019), and (2) tested for evidence for both dosage compensation and dosage balance in the Gila monster ZZ/ZW system.

**Figure 1.**
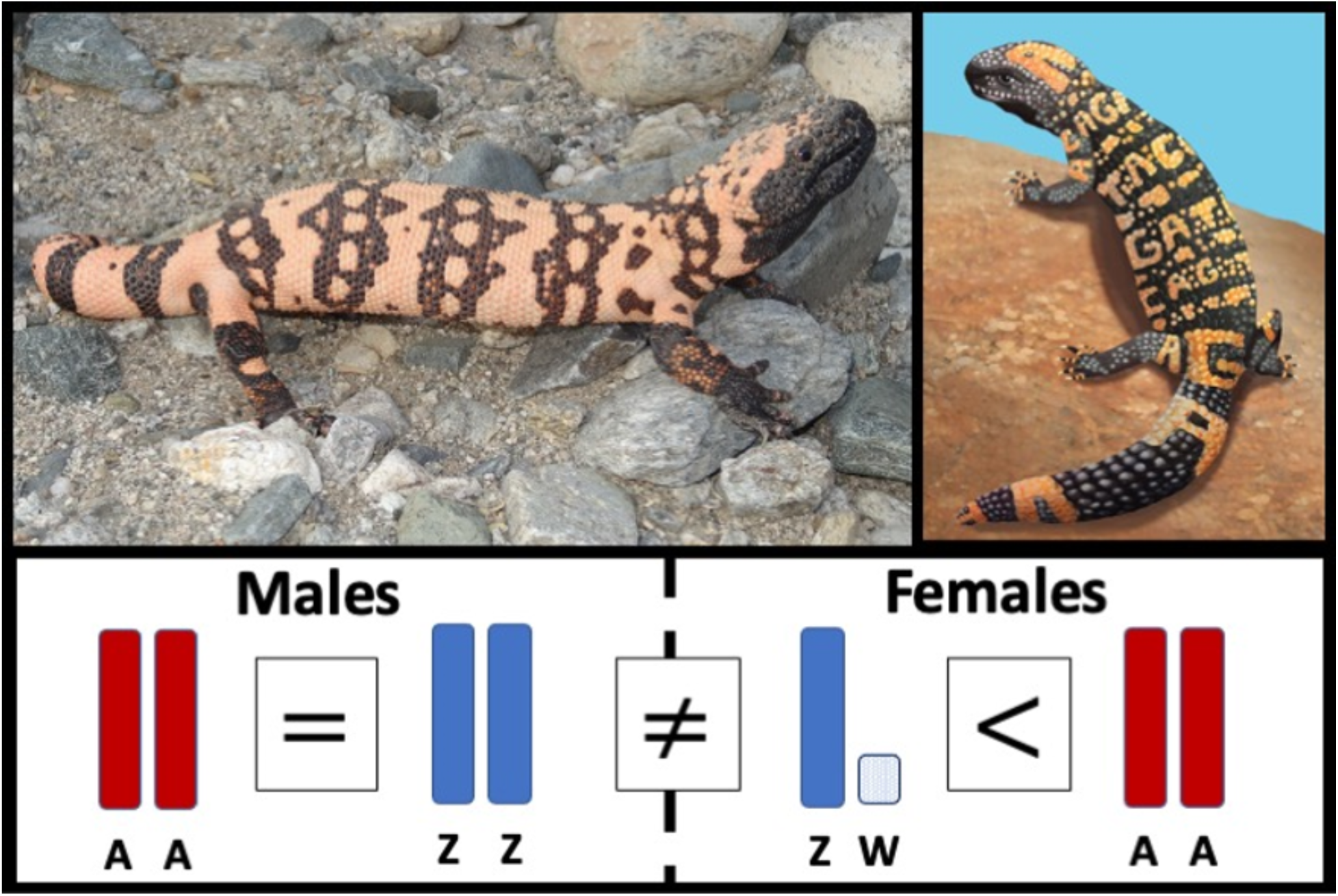
Graphical abstract. (Top Left) The Gila monster, *Heloderma suspectum*, with its distinctive black and orange pattern, is among the most iconic animals from the deserts of southwestern North America. (Top Right) The logo for this project, which started with a crowdfunding effort to assemble a reference genome in collaboration with 10X genomics. (Bottom) Using DNA and RNA data from six individuals (three males and three females), we investigated Gila monster sex chromosomes (ZW in females and ZZ in males) and their evolution, finding incomplete dosage balance between the sexes and a lack of dosage compensation.

## Materials and methods

### Samples and sequencing

We collected whole blood from the caudal vein near the tail base of six healthy, wild-born Gila monsters (*Heloderma suspectum*)—three males and three females. Blood samples for DNA sequencing were collected into 2 mL EDTA tubes (Supplemental Table S1), while blood samples for RNA sequencing were deposited into 1.5 mL tubes containing RNAlater (Table S2). All samples were immediately stored at –80°C.

We sent all samples to the Yale Center for Genome Analysis (YCGA) for extraction and sequencing. For whole-genome resequencing, samples were extracted following the YCGA’s standard protocol (Illumina TruSeq kit) and sequenced across two lanes of an Illumina HiSeq 4000 with 2×150bp paired-end sequencing. To minimize batch effects, we split males and females across the two lanes (i.e., 2 males and 1 female on lane 1, and 1 male and 2 females on lane 2). For RNA sequencing, RNA was extracted and prepared via the RiboZero protocol, after which samples were sequenced on a single lane of an Illumina HiSeq 4000 with 2×100bp paired-end sequencing.

### De novo reference genome assembly

We shipped whole blood from individual 10 (a ZZ male to improve our assembly of the Z chromosome) overnight on dry ice to 10× Genomics, where high molecular-weight genomic DNA was extracted and libraries were barcoded according to the Chromium Genome User Guide (details specified in (Weisenfeld et al., 2017). 10× Genomics generated approximately 140Gb of raw data on an Illumina HiSeq 2500, and used 115Gb of these data for the assembly generated with Supernova (Weisenfeld et al., 2017).

We calculated reference genome completeness and per-base quality statistics with kmers using merqury [v1.3] (Rhie et al., 2020). We further estimated genome completeness in a comparative framework using Benchmarking Universal Single-Copy Orthologs (BUSCO) [v5.1.2] (Simão et al., 2015), implemented on the gVolante web server [v2.0.0] (Nishimura et al., 2017)

### Genome annotation

Using Cactus (Armstrong et al., 2020), we aligned Gila monster to *Anolis carolinensis* (anoCar2) using garter snake (thaSir1), chicken (galGal5), and frog (xenTro9) as outgroups. The guide tree was ‘((Chicken:0.437442,(Anolis:0.247,(Gila:0.2,Garter_snake:0.2):0.1)1:0.2)1:0.172,Frog_X._tropic alis:0.347944).’ After alignment, Gila monster was annotated using the Comparative Annotation Toolkit (CAT; (Fiddes et al., 2018). To aid the annotation process, we aligned RNA-seq from 3 male Gila monsters and passed those alignments to CAT. We also used the RefSeq annotation of *A. carolinensis* as the source annotation set to lift to Gila monster. In addition, we predicted coding loci in all of the species simultaneously with the comparative annotation mode of Augustus (Nachtweide & Stanke, 2019).

### DNA alignment and variant calling

We assessed read quality with FastQC (Andrews, 2010) and MultiQC (Ewels et al., 2016). We used BBDuk (Bushnell et al., 2017) to remove adapter sequences and trim reads for quality (“`ktrim=r k=21 mink=11 hdist=2 tbo tpe qtrim=rl trimq=15 minlen=75 maq=20`”). Cleaned reads were mapped to our reference assembly with BWA MEM (H. Li, 2013) and duplicates were marked with SAMBLASTER (Faust & Hall, 2014), before using SAMtools (H. Li et al., 2009) to fix mates, and sort and index BAM files. We calculated basic BAM statistics using sambamba (Tarasov et al., 2015) (Supplemental Table S1).

For variant calling, we used GATK4 (Poplin et al., 2018). This multi-step process involves first calling variants in each sample separately with HaplotypeCaller (“`-ERC GVCF –– do-not-run-physical-phasing`”), then combining GVCF files from all six individuals with CombineGVCFs, and finally jointly calling variants across all samples with GenotypeGVCFs. To make variant calling more efficient, we divided the genome up into 25 segments of approximately equal size, running each of GATK4’s steps on each of these segments in parallel before using BCFtools (Danecek et al., 2021) to concatenate the resulting VCF files. Finally, we filtered variants for mapping quality (MQ >= 30), quality by depth (QD > 2), sample depth (FMT/DP >= 10), and GQ (FMT/GQ >= 30) with BCFtools. MQ and QD are site-wide measures, while DP and GQ filters were applied per sample.

### RNA mapping and quantification

We processed RNA reads from blood samples as described above for DNA reads, with the exception that we set “`minlen=60`” in BBDuk (Bushnell et al., 2017) because the RNA reads were shorter than those from DNA. We mapped reads using HiSat2 (Kim et al., 2019) with default parameters for paired-end reads before sorting reads with SAMtools (H. Li et al., 2009). We calculated basic BAM statistics using sambamba (Tarasov et al., 2015) (Supplemental Table S2). We next assembled transcripts using StringTie (Pertea et al., 2015) using a reference-based approach.

### Z chromosome scaffold identification

We identified candidate Z chromosome scaffolds using a two-step approach. First, we used the CHROM_STATS module in XYalign (Webster et al., 2019) with the “`--use-counts`” flag to gather mapped read counts per scaffold. As an approximation of depth of coverage, we divided the read count for each scaffold by the scaffold length and then took the mean of this value for males and females. We then calculated the mean female/male coverage per scaffold. While a number of scaffolds exhibited ratios substantially less than 1, as expected for a ZZ/ZW heterogametic system, values did not clearly separate into distinct Z and autosome clusters. When investigating other metrics across scaffolds, we discovered that five scaffolds, in addition to having some of the lowest female to male depth ratios across all scaffolds, also displayed extraordinarily high heterozygous rates in females (defined as the number of heterozygous sites over the number of non-reference sites). Of these scaffolds, four were longer than 500 kb and had greater than ten transcripts (scaffolds 157, 218, 304, and 398), and for the rest of the manuscript we treat these as candidate Z chromosome scaffolds. For our autosomal comparisons, we used the four largest scaffolds (0, 1, 2, 3), all of which had female:male depth ratios near 1 and exhibited female heterozygous rates that were neither close to 1 nor substantially higher than those of males.

Next, we scanned for pseudoautosomal regions (PARs) on the 4 putative Z scaffolds. For each Z scaffold, along with a representative autosomal scaffold (0), we obtained the log_2_ F/M ratio of DNA read depth in 5000 bp windows, calculated using XYalign (Webster et al., 2019). We used locally estimated scatterplot smoothing (LOESS) curves to visualize ratios by genomic location (Figure 2A), and manually inspected window depths in possible transition regions. While all Z scaffolds had lower overall read depth for females than males, consistent with expectations for female heterogamety, the first 1,750,000 bp of scaffold 304 showed balanced read depth in both sexes, suggesting a PAR (Figure 2A).

**Figure 2.**
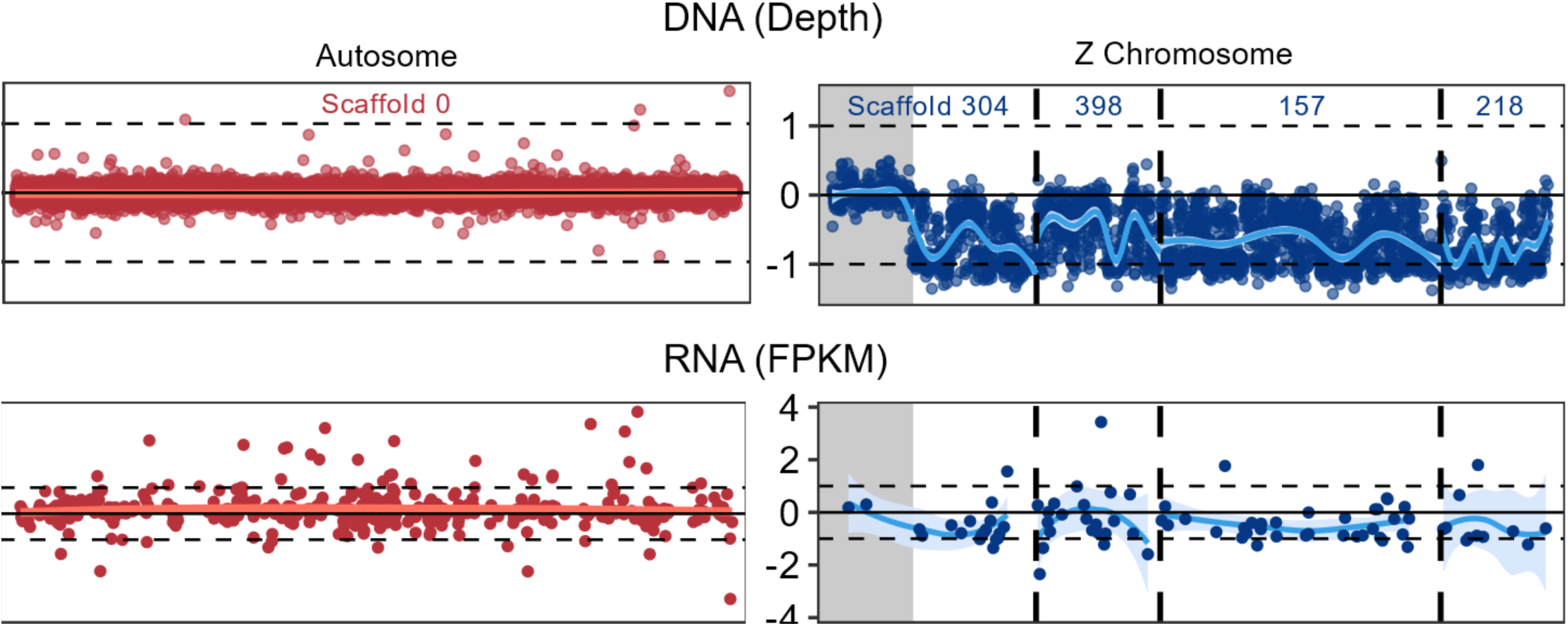
Identifying Gila monster Z-linked scaffolds. Log_2_-transformed ratios of Female:Male read depth from DNA (A) and Female:Male FPKM expression for RNA (B) for one autosomal scaffold (0; red) and the four putative Z chromosome scaffolds (304, 157, 218, and 398; blue). Log_2_ F/M ratios indicate higher (positive ratio) or lower (negative ratio) read depth and expression in females relative to males. LOESS curves show that read depth and transcript expression are balanced between the sexes for the autosome, but vary across scaffolds for the Z chromosome. We ordered scaffolds (separated by dashed lines) in the order in the *S. crocodilurus* genome (304, 389, 157), assigned by RagTag (see Materials and Methods), with the exception of 218, which was not mapped by RagTag and appended on the right end. Our hypothesized pseudoautosomal region is displayed in gray. Three female-biased DNA windows on the Z chromosome are not shown on the current plot due to making comparable axis between Z and autosomes, but were included in statistical analyses: two on scaffold 398 (ratios = 1.79 and 1.78), and one on scaffold 218 (ratio = 2.62).

For transcripts expressed in both sexes, we calculated mean expression values (FPKM) per transcript for each sex and visualized the log_2_(F/M) ratio of expression across the autosomal scaffold (0) and four Z scaffolds (Figure 2B). For most transcripts we observed lower expression in females compared to males (negative log_2_(F/M ratios)); however, some exhibited higher expression in females (positive log_2_(F/M ratios)), including 2 transcripts on the candidate PAR on scaffold 304 (Figure 2B).

### Synteny analyses

We used four different methods to identify syntenic regions between the Gila monster and other species. First, we used one-to-one orthologs identified by CAT during annotation to identify “ancestral” Gila monster autosomal and Z genes in chicken (*Gallus gallus*). As in Rovatsos et al. (2019), all orthologs of Z-linked genes in Gila monster are autosomal in chicken and located on chromosome 28.

Second, to assess synteny conservation, we employed bioinformatic synteny “painting” using a custom Perl script (*Gff2fasta.pl* modified from https://github.com/ISUgenomics/common_scripts), Biopython v1.73 (Cock et al., 2009), and conversion scripts from the CHROnicle package (v2015). We downloaded the genome FASTA and GFF annotation files of Komodo dragon (*Varanus komodoensis*; (Lind et al., 2019)) and then extracted and aggregated protein FASTA records using the modified Perl script *gff2fasta.pl*. Using the genome FASTA and a custom Python script longest_scaffolds.py, we identified the 24 longest scaffolds in the Komodo dragon genome. We extracted proteins from these scaffolds and six identified sex chromosome scaffolds from the protein FASTA records using a custom Python script *pull_id_match.py*. This was also performed for the 50 longest scaffolds and five putative sex chromosome scaffolds in Komodo dragon. SynChro computed conserved synteny blocks with delta=4, which requires four consecutive genes to match across species to be considered a syntenic block (Drillon et al., 2014).

Third, because synteny painting only successfully identified syntenic regions in the Komodo dragon genome for three of the four putative Z chromosome scaffolds in Gila monster, we used LastZ (Harris, 2007) to align the remaining scaffold to the entire Komodo dragon genome.

Finally, as we were finalizing this manuscript, the first chromosome-level genome of an anguimorph, the Chinese crocodile lizard (*Shinisaurus crocodilurus*), was published (Xie et al., 2022). To further resolve the order of scaffolds in Gila monster and Komodo dragon, we mapped these genomes to Chinese crocodile lizard using RagTag [v2.1.0] (Alonge et al., 2021) and visualized them using *pafr* [v0.0.2] (Supplemental Figure S2).

### Dosage compensation and dosage balance in Gila monster and chicken

In addition to the RNAseq data from three male and three female Gila monsters, we also included chicken (*Gallus gallus*) and green anole (*Anolis carolinensis*) as outgroups to approximate ancestral expression. We obtained publicly available RNAseq data from liver tissue for three male and female domestic chickens (Mullon et al., 2015); (NCBI BioProject PRJNA284655; females SRR2889291-3, males SRR2889295-7). Previous analyses confirm that patterns of dosage balance between autosomes and the Z chromosome are consistent across tissues in chicken (Zimmer et al., 2016) and are thus appropriate comparisons to the blood-derived RNAseq data from Gila monsters generated in this study. While we used chicken as the primary outgroup in our analyses because of its better annotation, we also confirmed results using green anole. For this species, we obtained publicly available RNAseq data from tail tissue (NCBI BioProjectPRJNA253971; (Hutchins et al., 2014; Rupp et al., 2017); (Rupp et al., 2017)). Though chicken and green anole differ in sex chromosome complement (ZZ/ZW and XX/XY, respectively), the Gila monster Z chromosome is syntenic with autosomal regions in both species. We processed the chicken and anole data using the same procedures as the Gila monster data (described above). We employed two statistical approaches to evaluate dosage balance and compensation: (1) nonparametric Mann-Whitney-Wilcoxon *U* tests—the most commonly used method for this problem in the literature—and (2) a linear modeling approach similar to that proposed by (Walters et al. 2015; Gu and Walters, 2017).

For our Mann-Whitney-Wilcoxon *U* analyses, we grouped genomic regions as follows: a Gila monster autosomal linkage group (syntenic with chicken chromosome 5) and Gila monster Z chromosome (scaffolds 157, 218, 304 without the putative PAR, and 398); a chicken autosome (chromosome 5), chicken chromosome 28 (syntenic with the Gila monster Z chromosome), and the chicken Z chromosome. To test for dosage balance (within species) and dosage compensation (between species), we compared F/M expression ratios in both chicken and Gila monster (Figure 3A) and relative expression for each region by sex (Figure 3B), respectively, using both frequentist (with Bonferroni corrections for multiple testing in each species) and Baysian Mann-Whitney-Wilcoxon tests using JASP [v0.16.2.0] (Figure 3; (JASP Team, 2022)). To alleviate confounding effects from potential microchromosome function (Perry et al., 2021), we dissected these expression data further by splitting autosomal genes out by their syntenic position in chicken showing the sex-specificity of the Gila monster Z relative to all other linkage groups (Supplemental Figure S1). For comparisons between chicken and Gila monster, we limited analyses to one-to-one orthologs identified by CAT during annotation (see *Genome Annotation* section above). For our Gila monster-green anole comparison, we separately identified one-to-one orthologs using OrthoFinder [v2.5.4] (Emms & Kelly, 2019).

**Figure 3:**
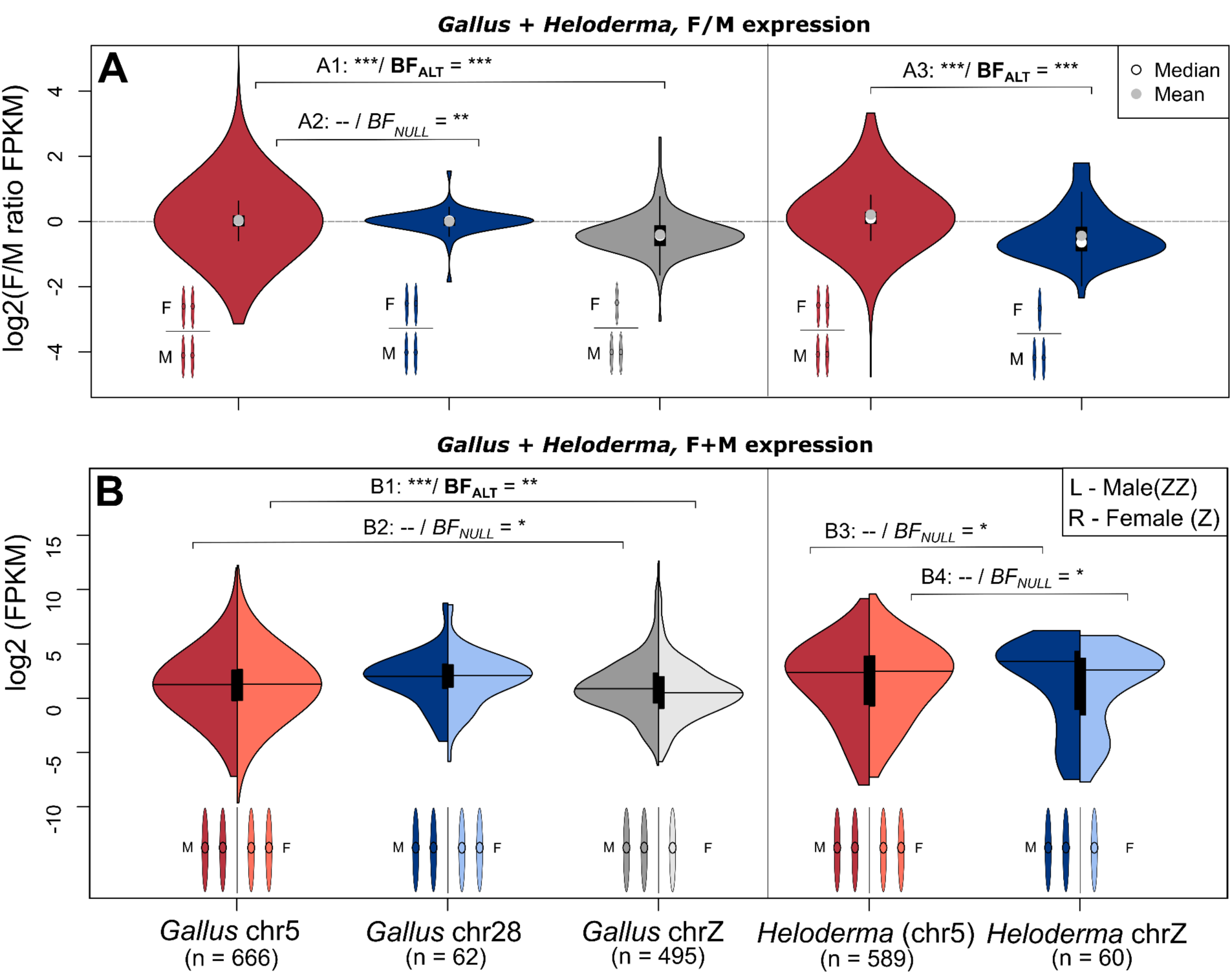
Complete dosage compensation without dosage balance in *Heloderma suspectum*. (A) Female/male FPKM transcript ratios across five genomic regions in *Gallus gallus* (chicken) and *Heloderma suspectum* (Gila monster): *Gallus* autosome (chr5), *Gallus* chr28, *Gallus* chrZ, *Heloderma* autosomes (syntenic with *Gallus* chr5), and *Heloderma* chrZ (scaffolds 157, 218, 304 (excluding putative pseudoautosomal region), and 398). Genomic regions *Gallus* chr5/*Heloderma* autosomal (shaded red) and *Gallus* chr28/*Heloderma* Z (shaded blue) are syntenic. (B) FPKM values separating male (left/darker) and female (right/lighter) violins for each of the five genomic regions. Statistics comparing each group are frequentist (Bonferroni corrected) and Bayesian Wilcoxon rank sum tests. P-values and Bayes Factors (Alt/Null) for each test (A1-B4) are reported in the main text. Support values summarized as: *** strong support, ** moderate support, * modest support.

We also tested for dosage balance and compensation using a linear modeling approach, as suggested by Walters and colleagues (Gu & Walters, 2017; Walters et al., 2015). It is possible that using means and ratios, as done with the Mann-Whitney-Wilcoxon *U* tests, masks important variation present in the data. In contrast, a linear mixed model (LMM) allows us to model individual and transcript variation in expression, along with our primary variables of interest. To this end, we fit sets of LMMs to test three conditions: (a) dosage balance, (b) dosage compensation using chicken as outgroup, and (c) dosage compensation using green anole as outgroup. We first normalized FPKM values using Ordered Quantile Normalization using the orderNorm transformation, the best supported normalization for the data estimated by the ‘bestNormalize’ package in R (Peterson, 2021; Peterson & Cavanaugh, 2020)). After transformation, we confirmed a normal distribution for the new data using the descdist function in the ‘fitdistrplus’ package (Delignette-Muller & Dutang, 2015). For all models, we included transcript ID and individual ID as random effects. In the dosage compensation models, we also included as a random effect the interaction between mean ancestral male and mean ancestral female expression for a given orthologous transcript, measured in the outgroup species. We reasoned that a difference in male and female Z chromosome expression present after controlling for this interaction would indicate divergence from relative ancestral expression and therefore a lack of dosage compensation (Gu & Walters, 2017; Walters et al., 2015). For each condition, we started with an intercept-only model and iteratively added sex, Z-linkage, and the interaction between the two as fixed effects. We conducted these analyses in R, using the package ‘lme4’ (Bates et al., 2015) to fit models, MuMIn (Bartoń, 2023) for model selection, and sjPlot (Lüdecke, 2023) for additional summary functions. We used AICc to determine the best supported model, treating models with ΔAICc of 2 or less as equally supported.

### Sexual selection and dosage balance

We tested the hypothesis that sexual selection might drive the lack of dosage balance across most ZZ/ZW systems following Mullon et al. (Mullon et al., 2015). Using the Gila monster RNAseq dataset described above, we first obtained read counts per sample per transcript using HTSeq (Putri et al., 2022). We then used edgeR (Robinson et al., 2010) to calculate the biological coefficient of variation (BCV), a measure of variability of expression, for each sex for each transcript and used the log2 of BCV for downstream analyses. Highly constrained expression is expected under strong purifying selection, while differences in variability between the sexes is a potential signature of a sex-bias in selection (Mullon et al., 2015; Romero et al., 2012).

Limiting our analyses to the non-PAR Z-linked genes expressed in both sexes that we identified in our dosage balance analyses described above, we tested two hypotheses: (1) selection should be more intense in males as a result of sexual selection stemming from greater variance in reproductive success among males than females, and because of this, (2) dosage balance on the Z chromosome should occur on a gene-by-gene basis in genes under strong selection in either females or both sexes. We tested the first hypothesis by comparing BCV between the two sexes with a Wilcoxon signed-rank test in R. Because male and female expression tend to be correlated, we ran a PCA to project variability across two orthogonal axes

(Mullon et al., 2015). Like Mullon et al. (2015), we found that PC1 corresponded to the intensity of sexually concordant selection, while PC2 represented sex-bias, with greater values indicating a stronger male bias. We used these two variables and their interaction as predictors in a linear model, with log2(Female FPKM / Male FPKM) as our outcome.

## Results and Discussion

### Gila monster draft genome assembly

We sequenced and assembled a draft genome assembly for the Gila monster (*Heloderma suspectum*) using DNA collected from a wild-born male (ZZ) from Arizona (USA) housed at Arizona State University. The final haploid genome assembly was 2.56Gb in total length, with a scaffold N50 of 7.86Mb and a contig N50 of 35.49Kb (Supplemental Tables S3 and S4). Interestingly, this genome assembly was the largest of any available available anguimorph genome, 70%, 25%, and 12% larger than Komodo dragon (*V. komodoensis*), Chinese crocodile lizard (*S. crocodilurus*), and beaded lizard (*H. charlesbogerti*) assemblies, respectively (Pinto et al., 2023). The assembled Gila monster genome was 97.2% complete— calculated using kmers—with an average per-base error rate of 7.15316 x 10^-05^ (i.e. <1 error per 10kb). When estimating genomic completeness in a comparative framework using BUSCO [v5.1.2] (Simão et al., 2015), querying two databases (Sauropsida and Core Vertebrate Genes [CVG] (Hara et al., 2015), we found that our assembly maintains a >90% completeness score. For the Sauropsida database of 7,480 genes, the assembly contains 90.9% complete orthologs, with 1.2% duplicated, 3.4% fragmented, and 5.7% missing. For the CVG database of 233 genes, the assembly contains 94.8% complete orthologs with 0% duplicates, 3.0% fragmented, and 2.2% missing. Thus, the Gila monster genome is largely complete and accurate.

During genome annotation, CAT identified 15,721 genes in the assembly. 15,129 of these were identified as orthologs of genes in the RefSeq annotation of green anole (18,595 genes). The remaining 1,007 genes came from comparative Augustus predictions (Stanke et al.,

2006). Of those 1,007 predictions, there were 131 putatively novel loci, while 617 were predicted to be paralogs, and thus candidates for gene family expansion events. 37 genes had evidence of being split into multiple locations on a single contig, and a further 380 genes had evidence of being split across multiple contigs. To examine the completeness of genome annotation, we again used BUSCO [v5.1.2] (Simão et al., 2015). If the annotation captured most genes present in the genome assembly, the BUSCO scores should be comparable to the unannotated assembly. For the Sauropsida database of 7,480 genes, the annotations contain 75.9% complete orthologs, with 2.5% duplicated, 6.9% fragmented, and 17.2% missing. For the CVG database of 233 genes, the assembly contains 85.8% complete orthologs with 3.4% duplicates, 7.3% fragmented, and 6.9% missing. Both evaluations of annotation completeness presented much lower scores than that of the full assembly, leaving room for future improvement of the genome annotation.

### Identifying sex chromosome scaffolds in the Gila monster

We identified four putative Z-linked scaffolds, greater than 500kb in length, within the Gila monster genome assembly using mean F/M read depth (Table 1). These four scaffolds (157, 218, 304, and 398) also exhibited an extreme excess of heterozygous sites in females relative to males (Supplemental Table S5). While male heterozygosity on these scaffolds overlapped with autosomal heterozygosity, the average number of heterozygous sites on these scaffolds in females ranged from 13-35 times that of males. Though genetic diversity on the sex chromosomes can be affected by a number of processes (Webster & Wilson Sayres, 2016; Wilson Sayres, 2018), it is unlikely to explain these results for three reasons. First, since Z chromosomes are inherited by both sexes, estimates of diversity should not differ between males and females. Second, Z/A ratios have a theoretical maximum less than 1.2 (Charlesworth, 2009; Corl & Ellegren, 2012), an order of magnitude less than the values observed here. Third, outside of pseudoautosomal regions, females should not have any heterozygous sites because they possess a single Z. Instead, we suggest that this is due to reads from the female W chromosomes mismapping to the Z (Pinto et al., 2022; Schield et al., 2019; Webster et al., 2019) because there is no W chromosome in the assembly. To explore this further, we called variants in the RNAseq data and observed similar heterozygous rates in males and females (Supplemental Table S5). It is unclear why the use of RNA would reduce heterozygous rates to more realistic values. However, the similar heterozygous rates in RNA between males and females, the latter of which should lack heterozygous sites, is consistent with mismapping between gametologs.

**Table 1.**
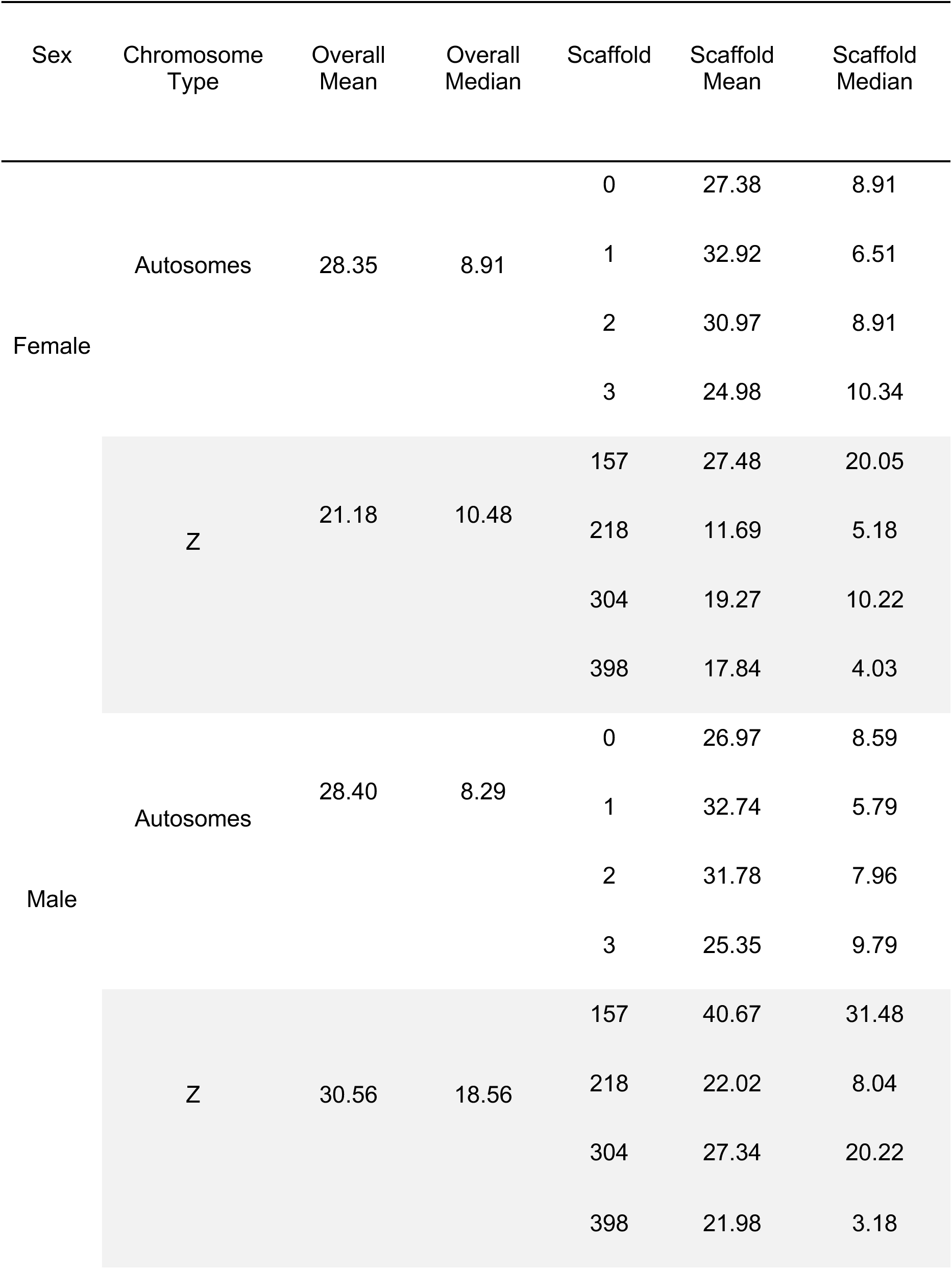
Summary statistics for each scaffold used for dosage analyses. Means and medians for 3 males and 3 females are presented as FPKM values after filtering out unexpressed transcripts (FPKM = 0 in either sexes). After filtering, 1,099 and 86 transcripts remained across the autosomal and Z scaffolds, respectively. Transcripts from the putative PAR on Z scaffold 304 were removed prior to the above calculations.

To better understand the sex-specificity of these putative Z scaffolds, we examined F/M read depth in 5,000bp windows (DNA) and per-transcript F/M expression in FPKM (RNA) along the Z chromosome, relative to an autosome (scaffold 0; Figure 2). Autosomal transcripts varied more than Z transcripts in their F/M expression ratios, though LOESS curves indicated relatively balanced autosomal expression. The Z scaffolds, on the other hand, displayed consistently negative (male-biased) expression ratios regardless of scaffold location. However, in a 1.75Mb region at the beginning of scaffold 304, read depth and expression ratios matched those of the autosome (Figure 2), suggesting a pseuo-autosomal region (PAR). Interestingly, scaffold 304 mapped most proximally, relative to other sex-linked scaffolds, to Chinese crocodile lizard chromosome 7 (Supplemental Figure S2). We also highlight one scaffold, 674, which was excluded because of its length (<500kb) and few annotated transcripts (4), but had a low mean female:male read depth and high heterozygosity in females (Supplemental Table S5). We also found that one gene on this scaffold also maps to chicken chromosome 28 suggesting it too is likely\ part of the Z chromosome linkage group in Gila monster.

### Synteny of Z chromosome scaffolds in Gila monster

Synteny painting analyses with SynChro (Drillon et al., 2014) revealed that three of the four sex-linked scaffolds in Gila monster are largely syntenic with three corresponding scaffolds in Komodo dragon (Figure 4). Further, mapping these scaffolds to the Chinese crocodile lizard genome showed that these sex-linked Gila monster scaffolds (304, 398, and 157) and Komodo dragon scaffolds (SJPD01000091.1, SJPD01000092.1, and SJPD0100101.1) all co-localize to the distal region of chromosome 7 (Supplemental Figure S2). Because the fourth Z scaffold, scaffold 218, did not map with SynChro or RagTag, we used LastZ (Harris, 2007) to align it with the entire Komodo dragon assembly. The top alignment hit in Komodo dragon was also SJPD01000092.1 (22,510 bp aligned). Therefore, across reptiles, the sex chromosome linkage group in Gila monster and Komodo dragon correspond to chromosome 7 in Chinese crocodile lizard (Anguimorpha), LgB in green anole (Iguania), and chromosome 28 in chicken (Aves).

**Figure 4.**
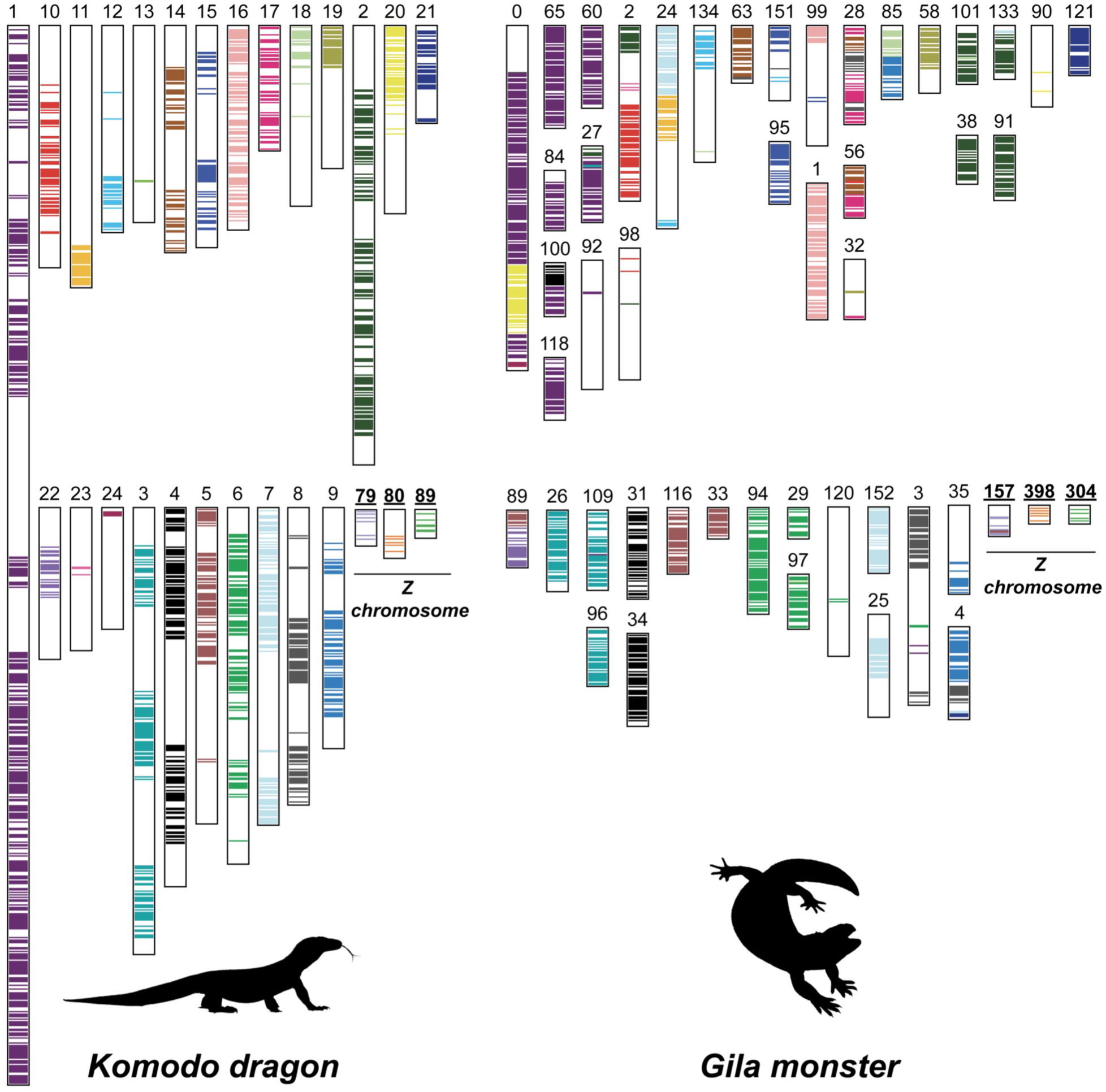
Genome homology between Komodo dragon and Gila monster. For each putative sex chromosome scaffold and the 24 longest scaffolds, we show their position in the Komodo dragon (*V. komodoensis*) genome from Lind et al. (2019) and the Gila monster *(H. suspectum*) genome. Komodo dragon scaffold numbers correspond to their names and annotations on Figshare (https://doi.org/10.6084/m9.figshare.7949483.v2), but the three sex chromosome scaffolds are known as 79:SJPD01000091.1, 80:SJPD01000092.1, and 89:SJPD01000101.1 on Ensembl [v105.1].

**Figure 5.**
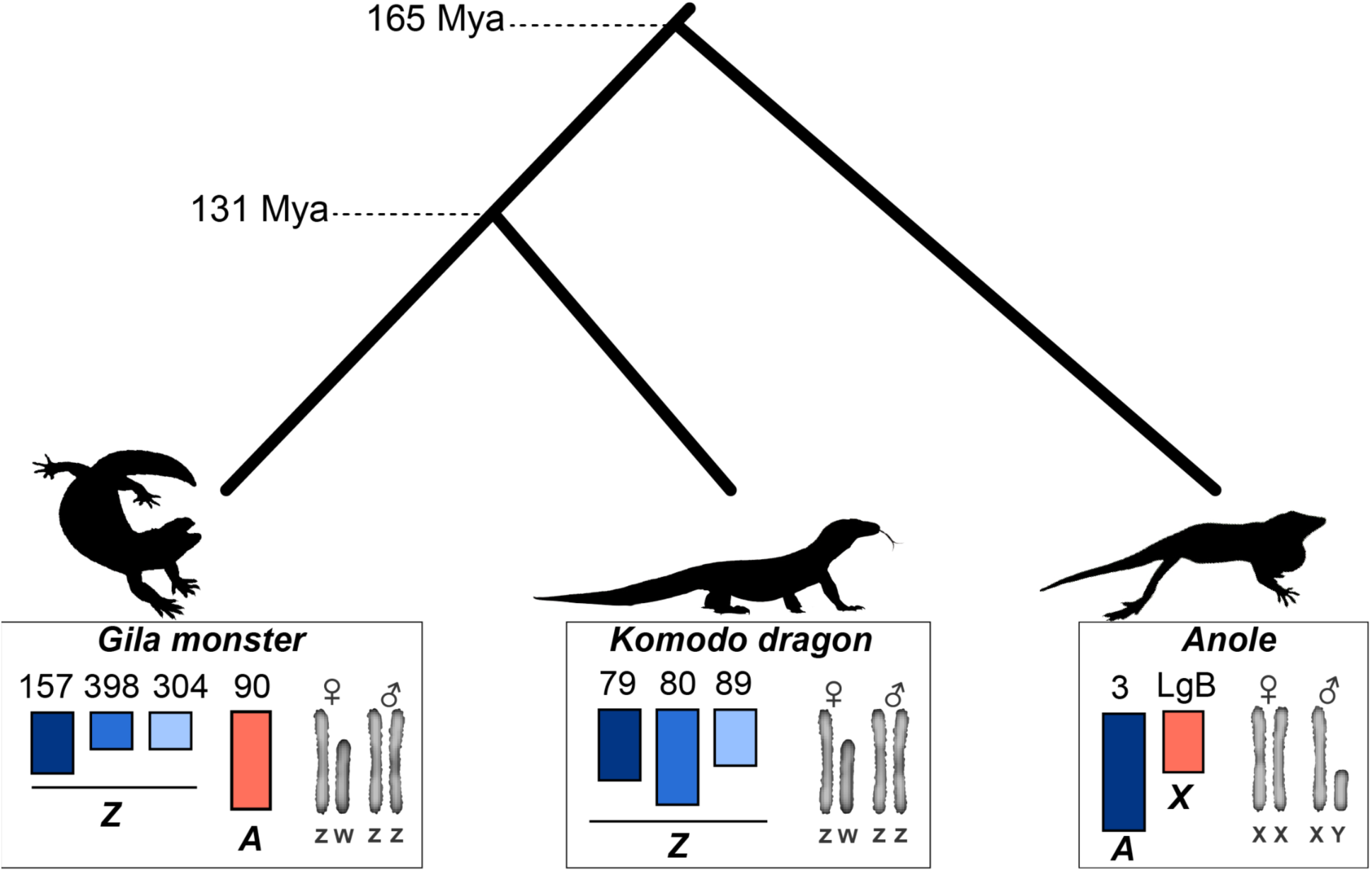
Gila monster sex chromosome sequence similarity with Komodo dragon and green anole. Here we show the phylogenetic tree of the relationship among Gila monster (*H. suspectum*), Komodo dragon (*V. komodoensis*), and green anole (*A. carolinensis*), with approximate divergence times from TimeTree.org. The Komodo dragon scaffold numbers correspond to their names and annotations on Figshare (https://doi.org/10.6084/m9.figshare.7949483.v2), but the three sex chromosome scaffolds are known as 79:SJPD01000091.1, 80:SJPD01000092.1, and 89:SJPD01000101.1 on Ensembl [v105.1].

### Insight into sex determination mechanisms

We further investigated the functional annotations of genes in the putative Gila monster Z scaffolds. The homologous linkage group in chicken, Gg28, contains the anti-Müllerian hormone (*Amh*) gene, a gene involved in testis differentiation known to act as a master sex determining gene in multiple groups of fishes (M. Li et al., 2015; Myosho et al., 2015; Pan et al., 2017). *Amh* is retained on this linkage group in Gila monster and, in blood tissue, is expressed twice as high in males than females (F/M ratio = 0.513). This linkage group, including *Amh*, is also found on a sex chromosome linkage group in monotremes (Kratochvíl et al., 2021).

However, as there are few other potential candidate genes presently assembled on this linkage group, more evidence, (at a minimum) a more complete list of Z-linked genes, is needed to explicitly implicate *Amh* as a candidate primary sex determining gene in Gila monster and other anguimorphs.

### Lack of dosage balance with incomplete dosage compensation in Gila monster

To initially test for dosage balance in Gila monster, we isolated autosomal and sex-linked gene expression data in chicken and Gila monster and used Mann-Whitney-Wilcoxon tests in frequentist and Baysian frameworks. Frequentist statistics are most commonly used in this scenario, however, Bayesian inference can help provide a more nuanced picture (i.e. show support for the *NULL* and *ALT* hypothesis with varying thresholds, where from Bayes Factors (BF)>30 are considered strong support to 10>BF>1 are considered modest support). The number of universally-expressed transcripts (expressed in both sexes) present on a representative autosome (Gg5) in chicken and Gila monster were 666 and 589 transcripts, respectively. We filtered to include only expressed transcripts with 1:1 orthologs on the syntenic chicken chromosome 28 and Gila monster Z leaving 62 and 60 transcripts, respectively (Supplemental Table S6). Lastly, there were 495 transcripts expressed on the chicken Z chromosome. We used these data to test for dosage balance between the sexes and found that F/M gene expression was lower on the Z chromosome in both chicken and Gila monster (Figure 3A; A1 p-value = 2.73 x 10^-57^ & BF_ALT_ = 2.2 x 10^8^, and A3 p-value = 1.5 x 10^-13^ & BF_ALT_ = 257, respectively). This pattern was previously identified in chicken and replicated here for comparative purposes (Supplemental Figure S1; Ellegren et al., 2007; Itoh et al., 2007). The log_2_ ratios of Z chromosome genes in Gila monster (mean = –0.44, median = –0.64) are higher than what is expected with a complete lack of dosage balance (i.e. approximately –1.0) (Schield et al., 2019). The linear modeling approach recapitulated the Mann-Whitney *U* results and also identified a lack of dosage balance in Gila monster (Table 2a; Table 3a). The full model, which included sex, Z-linkage, and their interaction as fixed effects, performed best (ΔAICc ≥ 161.11) and was the only model better than the null model (Table 2a). In this model, the interaction between sex and Z-linkage was the only significant term (male * Z-linked β = 0.25), consistent with higher expression on the Z in males than females. Thus, the lower F/M expression on sex chromosomes, relative to autosomes, indicates a state of incomplete dosage balance between the sexes (Gu & Walters, 2017).

**Table 2.** Results of model selection. The outcome variable was expression (normalized FPKM) for all models. (A) “Gila only” models included individual (1 | individual) and transcript (1 | transcript) IDs as random effects. (B and C) Both sets of models controlling for ancestral expression included those variables along with an interaction between male and female expression in the outgroup (1 | anc_male:anc_female) as an additional random effect. Within each cluster, models are ordered by ΔAICc values, with the best model listed first.

| Model | Intercept | $\Delta AICc$ | Weight |
| --- | --- | --- | --- |
| *a) Gila only - Dosage Balance* |  |  |  |
| Sex + Z_linkage + Sex * Z_linkage | 0.0035 | 0 | 1 |
| Null | -0.0124 | 161.11 | 0 |
| Sex | -0.0033 | 162.67 | 0 |
| Z_linkage | -0.0144 | 163.05 | 0 |
| Sex + Z_linkage | -0.0053 | 164.61 | 0 |
| *b) Gila controlling for ancestral expression (chicken) - Dosage Compensation* |  |  |  |
| Sex + Z_linkage + Sex * Z_linkage | 0.0387 | 0 | 1 |
| Null | 0.0151 | 229.35 | 0 |
| Z_linkage | 0.0188 | 232.35 | 0 |
| Sex | 0.0236 | 232.85 | 0 |
| Sex + Z_linkage | 0.0273 | 233.85 | 0 |
| *c) Gila controlling for ancestral expression (anole) - Dosage Compensation* |  |  |  |
| Sex + Z_linkage + Sex * Z_linkage | -0.00136 | 0 | 1 |
| Null | -0.03125 | 50.94 | 0 |
| Sex | -0.01508 | 53.12 | 0 |
| Z_linkage | -0.02215 | 53.54 | 0 |
| Sex + Z_linkage | -0.00602 | 53.72 | 0 |

**Table 3.** Results of best linear mixed models evaluating (a) dosage balance and (b and c) dosage compensation. In all three cases, the full model had the lowest AICc value and was thus considered to be the best model (Table 2).

| Fixed effect | Estimate | Std. error | P |
| --- | --- | --- | --- |
| *a ) Gila only - Dosage Balance - Full Model* |  |  |  |
| Intercept | 0 | 0.04 | 0.919 |
| Sex (Male) | -0.04 | 0.03 | 0.178 |
| Z-linkage (Z) | -0.10 | 0.11 | 0.379 |
| Sex (Male) x Z-linkage (Z) | 0.25 | 0.02 | <0.001*** |
| *b) Gila controlling for ancestral expression (chicken) - Dosage Compensation - Full Model* |  |  |  |
| Intercept | 0.04 | 0.04 | 0.335 |
| Sex (Male) | -0.04 | 0.02 | 0.085 |
| Z-linkage (Z) | -0.2 | 0.14 | 0.142 |
| Sex (Male) x Z-linkage (Z) | 0.32 | 0.02 | <0.001*** |
| *c) Gila controlling for ancestral expression (anole) - Dosage Compensation - Full Model* |  |  |  |
| Intercept | 0 | 0.05 | 0.997 |
| Sex (Male) | -0.04 | 0.02 | 0.057 |
| Z-linkage (Z) | -0.47 | 0.26 | 0.074 |
| Sex (Male) x Z-linkage (Z) | 0.32 | 0.04 | <0.001*** |

We first attempted to diagnose dosage compensation status in the Gila monster using Mann-Whitney *U* tests, the most common approach for this problem (Gu & Walters, 2017), and the combination of (1) a Z-to-autosome comparison within Gila monster and (2) an ancestral state comparison proxied by comparing the syntenic linkage group (Gg28) to other autosomes in chicken. We found no significant differences in within-sex Z-to-autosome expression between male and female Gila monster (Figure 3B; B3: males, p-value = 0.851 & BF_NULL_ = 7.87, and B4: females, p-value = 0.157 & BF_NULL_ = 4.69), a pattern distinct from chicken, which lacks global dosage compensation (Figure 3B; B1: females, p-value = 3.53 x 10^-7^ & BF_ALT_ = 13.86, and B2: males, p-value = 0.027 & BF_NULL_ = 4.48). Further, we found no sex-biased expression patterns in our ancestral proxy, chicken chromosome 28, relative to autosomes (Figure 3A; p-value = 0.783 & BF_NULL_ = 12.29). Equal expression between the Z chromosome and autosomes for both males (ZZ) and females (Z) suggested complete dosage compensation on the Z chromosome in Gila monster. There is little evidence for this pattern (Type IV: complete dosage compensation without balance) in nature (Gu & Walters, 2017), which suggested that our results might be driven by a statistical artifact. Possible explanations include a small sample size (only ∼60 Z-linked transcripts in Gila monster with orthologs in chicken), variance in expression among transcripts, and differences in expression among individuals. Thus, traditional statistical examinations of dosage compensation may have been underpowered to resolve the dosage compensation status in this system.

As with dosage balance above, we reanalyzed these data in a linear mixed model (LMM) framework as proposed by Walters and colleagues (Gu & Walters, 2017; Walters et al., 2015), in which we can account for variation among transcripts and individuals. Using chicken as our outgroup and ancestral proxy, our best model was the full model (ΔAICc ≥ 229.35), in which the interaction between sex and Z-linkage was the only significant term (male * Z-linked β = 0.32; Table 2b; Table 3b; Figure S3). Replacing chicken with green anole produced the same qualitative results, confirming that choice of outgroup did not affect our analyses (Table 2c; Table 3c). These results are consistent with incomplete dosage compensation, as Gila monster Z chromosome expression in females remained lower than that of males while controlling for ancestral expression. As this linear model approach both replicated results obtained by our traditional statistical approaches and extended beyond their limitations, we strongly recommend this approach for future studies of dosage compensation.

We therefore infer that Gila monsters possess a ZZ/ZW system characterized by a lack of dosage balance and incomplete dosage compensation. This pattern (Type III in Gu & Walters, 2017) has been observed in almost every ZZ/ZW system that has been studied, the major exception being Lepidoptera (Gu & Walters, 2017). Previous work has shown that dosage compensation in ZZ/ZW systems can occur on a gene-by-gene basis (Graves, 2016a; Gu & Walters, 2017). Our data suggest that this is likely the case for the Gila monster as well, as average female Z expression was greater than half that of males and we observed substantial gene-by-gene variation in relative female Z expression, including multiple transcripts with greater female than male expression (Figure 2b).

Why global dosage compensation would be more important in male heterogametic than female heterogametic systems remains unclear (Chen et al., 2020; Gu & Walters, 2017; Naurin et al., 2010). Given that Gila monster sex chromosomes date back to at least the early Cretaceous or late Jurassic (>115 mya), they stand among some of the the oldest known vertebrate sex chromosomes and the extant lack of global dosage compensation cannot be explained by ‘a lack of time for it to have evolved’—as if dosage compensation were an inevitable outcome of sex chromosome evolution. Another explanation that has been proposed involves sexual selection, whereby greater reproductive skew in males could lead to more intense selection on expression (Mullon et al. 2015). Models suggest that this could lead to rapid global dosage compensation in male heterogametic systems and slower, more mosaic compensation in female heterogametic systems, where only genes under strong selection in females evolving compensation (Mullon et al. 2015). We explored two hypotheses related to this explanation in the Gila monster. In comparing selection (using BCV, a measure of variability in expression) between the sexes, we found evidence of more intense selection on expression in males than females (male mean = –1.53, female mean = 0.87; Wilcoxon signed rank test p <

1.148 x 10^-15^, V=0), a result also observed in chickens (Mullon et al. 2015). However, in contrast to chickens (Mullon et al. 2015), we found no effect of sexually concordant selection or more intense female selection on dosage balance (Table S7; Supplemental Figure S4). Thus, the lack of global dosage on the ancient Gila monster Z chromosome remains a mystery and an important avenue of future research.

## Conclusions

Here, we presented the draft genome assembly of the Gila monster, *H. suspectum*, alongside DNA re-sequencing and RNAseq data for multiple male and female individuals. We identified four scaffolds (>500kb) with male-biased patterns of read mapping and gene expression (Figure 2). We confirmed that these scaffolds are syntenic with the Komodo dragon (*V. komodoensis*) Z chromosome and chicken (*G. gallus*) chromosome 28 (Gg28), as shown previously shown (Rovatsos et al., 2019).

We found a patterns of expression consistent with incomplete dosage balance between the male and female Z chromosomes and autosomes (consistent with previous data from varanids; Rovatsos et al., 2019) and incomplete dosage compensation between the Z chromosome and their ancestral autosomal pair (Figure 3). This pattern has been observed in most other ZZ/ZW systems studied to date and may represent a more general pattern for ZZ/ZW systems (Gu & Walters, 2017). Our assembly of the Gila monster genome contained relatively few Z-linked genes and we could not resolve dosage compensation with the nonparametric tests typically used in these analyses. However, a linear modeling approach, in which we were able to account for variation among transcripts and individuals, allowed us to successfully infer the presence of incomplete dosage compensation. We suggest that other researchers consider this approach for similar analyses. Taken together, this work adds to our understanding of sex chromosome evolution in squamates and more generally.

## Data and code availability

Data generated in this study, including the *Heloderma suspectum* genome assembly, have been deposited to the NCBI SRA under Bioproject PRJNA420754. The versions of genome assembly and annotation files used in these analyses have been deposited in Zenodo (Webster et al., 2022). Steps for processing and analyzing RNA and DNA sequencing were built into a Snakemake (Mölder et al., 2021) pipeline, with all software managed via Bioconda (Grüning et al., 2018) in a Conda environment. All code for this pipeline and environment (including software versions) is available on Github: https://github.com/thw17/Gila_sex_chroms. Code used in the synteny painting analyses is available at https://github.com/mmoral31/Gila_Macrosynteny_Pipeline.

## Acknowledgements

We are tremendously grateful for a collaboration with 10XGenomics, who contributed the SuperNova genome assembly, that allowed us to extend the research to include analysis of DNA and RNA from six individuals in addition to assembly of a reference genome. We would like to thank George (PJ) Perry for advice in getting our crowdfunding campaign started, Daniel Beck, Carlos Infante, Tony Gamble, and Taylor Edwards for endorsing our project, members of the Wilson lab who helped promote the project (Shawn Rupp, Kimberly Olney, George A. Brusch IV, Pooja Narang, Sarah Brotman, Daniel Cotter, Ephrance Peninah Kalungi, and Valeria Valverde-Vesling), Experiment.com for working with us to make a beautiful campaign, and all of our 173 named and anonymous backers, without whom, this research would not have happened: St. Augustine Alligator Farm Zoological Park, Susan Allardyce, Manuel Ares Jr., Karen S. Ashe, Oroszlany Balazs, Nick Banovich, Joan Baron, Susan Bates, Susan Bello, Colleen Benton, Aatish Bhatia, Tim Blankenship, Computer Bob, Amy Boddy, Lawrence Mark Brotman, Sarah Brotman, Dave Bruce, Michael Cardwell, Gloria Carr, Bob Catt, KB Choi, Matthew Cichocki, Blair Costelloe, Richard Cotter, Daniel Cotter, Matt Cover, Tracey Depellegrin, Laura DePriest, Isabella T. Dorr, Joshua Drew, Peg Duthie, Andy Eguiluz, Anders Eklund, Noah Fahlgren, Nico Franz, Genome Galaxy, Tony Gamble, Jacquelyn Gill, Jose Gonzalez, Monica Gonzalez, Bastian Greshake, Aditya Gune, Penny Gwynne, Carl Hannah, Chad Harland, Susan Hayden, John Heidecker, Joshua Herr, Katie Hinde, Alexander Hirschi, Kathleen Holladay, Paul A. Hoskisson, TC Houston, Courtney Hoyt, Jacque and Peter Hoyt, Brandon Hurr, Elizabeth Hutchins, Chris Hyde, Karen James, Domino Joyce, Ephrance Peninah Kalungi, Charles Kazilek, Caroline Keroack, Jacob G. Kirkland, Kristine Klewin, Michelle Kline, Christine Kovach, Robin K. Kurtzner, Anja Landsmann, Denny Luan, Dylan MacKay, Mitch Magee, Michelle Marshall, Stephanie Marson, Kirk Michael Maxey, Lisa McCann, Liam McIver, Scott McMillin, Lem Meade, Donna DeVore Metler, Laura Meyer, Jennifer Meyer, Mark F. Miller, Aaron Minoo, Asia Murphy, Radu Nan, Pooja Narang, Randolph Nesse, Nick Nevid, Sue O’Brien, Kimberly Olney, Richard Otwell, Ty Park, Charlyn Partridge, John-Alan Pascoe, William Paulus, Lidia Peon, Ethan Perlstein, George (PJ) Perry, Anali Perry, Barret Phillips, Tanya Phung, Timothee Poisot, Dianne Elizabeth Price, Pascal Pucholt, Ellen Quillen, Michael Radis, Jennifer Raff, Linda Raish, Kate Ray, Ryan Remus, Roger Repp, Lila Robinwood, Matthew Rolfes, Phillip James Rose, Renee Rosier, Christie Rowe, Eric Samorodnitsky, Shantanu Sane, Thomas Schmutzer, Mark Seward, Marion Shadbolt, Ralph Shepstone, Sheetal Shetty, Vinay Singh, Anne Sitamun, Christopher Irwin Smith, Mark Soufleris, Bonnie Stewart, Anne Stone, Emily Taylor, Roy Toft, Barney Tomberlin, Christina Tran, Todd Trowbridge, McKenna Valverde-Vesling, Valeria Valverde-Vesling, Arvind Varsani, Craig Vesling, Wendy Vesling, Vesling Consulting Inc, Mona Vijay, Eric Damon Walters, John Webster, Bo Webster, Paula And Ralph Webster, Doris and Harvey Webster, Michell Werner, Jason Williams, Dottie Wilmore, Deanda Farr Wilson, George Wilson, David Winter, Cindy Wu, and Jeremy B. Yoder. This work was also supported by the National Institute of General Medical Sciences (NIGMS) of the National Institutes of Health (NIH) grant R35GM124827 to MAW. We further thank Arizona State Research Computing and the Center for High Performance Computing at the University of Utah for computational support and resources. Finally, we are grateful to Brian Codding, Thomas Kraft, and Alan Rogers for helpful statistical discussions and the Primate Evolution and Genomics Lab at the University of Utah for comments on the manuscript.

## Supplementary Materials

|  |  |
| --- | --- |
| Supplemental Table S1: DNA Resequencing Sequencing Statistics for Gila Monster. |  |
| Supplemental Table S2: RNA Sequencing Statistics for Gila Monster. |  |
| Supplemental Table S3: Gila monster (*Heloderma suspectum*, male #10) Genome Statistics, part 1. |  |
| Supplemental Table S4: Gila monster (*Heloderma suspectum*, male #10) genome statistics, part 2. |  |
| Supplemental Table S5. Per-scaffold depth, expression, and genotype statistics. | .xlsx file |
| Supplemental Table S6: *Heloderma suspectum* and *Gallus gallus* gene expression on Gg28 | .xlsx file |
| Supplemental Table S7. Results of linear model testing the effect of selection intensity on dosage balance. |  |
| Supplemental Figure S1: F/M gene expression grouped by chicken chromosomes. |  |
| Supplemental Figure S2: *Heloderma suspectum* and *Varanus komodoensis* genomes aligned to the recently published *Shinisaurus crocodilurus* genome. |  |
| Supplemental Figure S3: Marginal effects of the interaction between sex (ZZ: male; ZW: female) and chromosome type (autosome vs. Z chromosome) on expression in Gila monster. |  |
| Supplemental Figure S4. Selection and dosage balance in Gila monster. |  |

**Supplemental Table S1:**
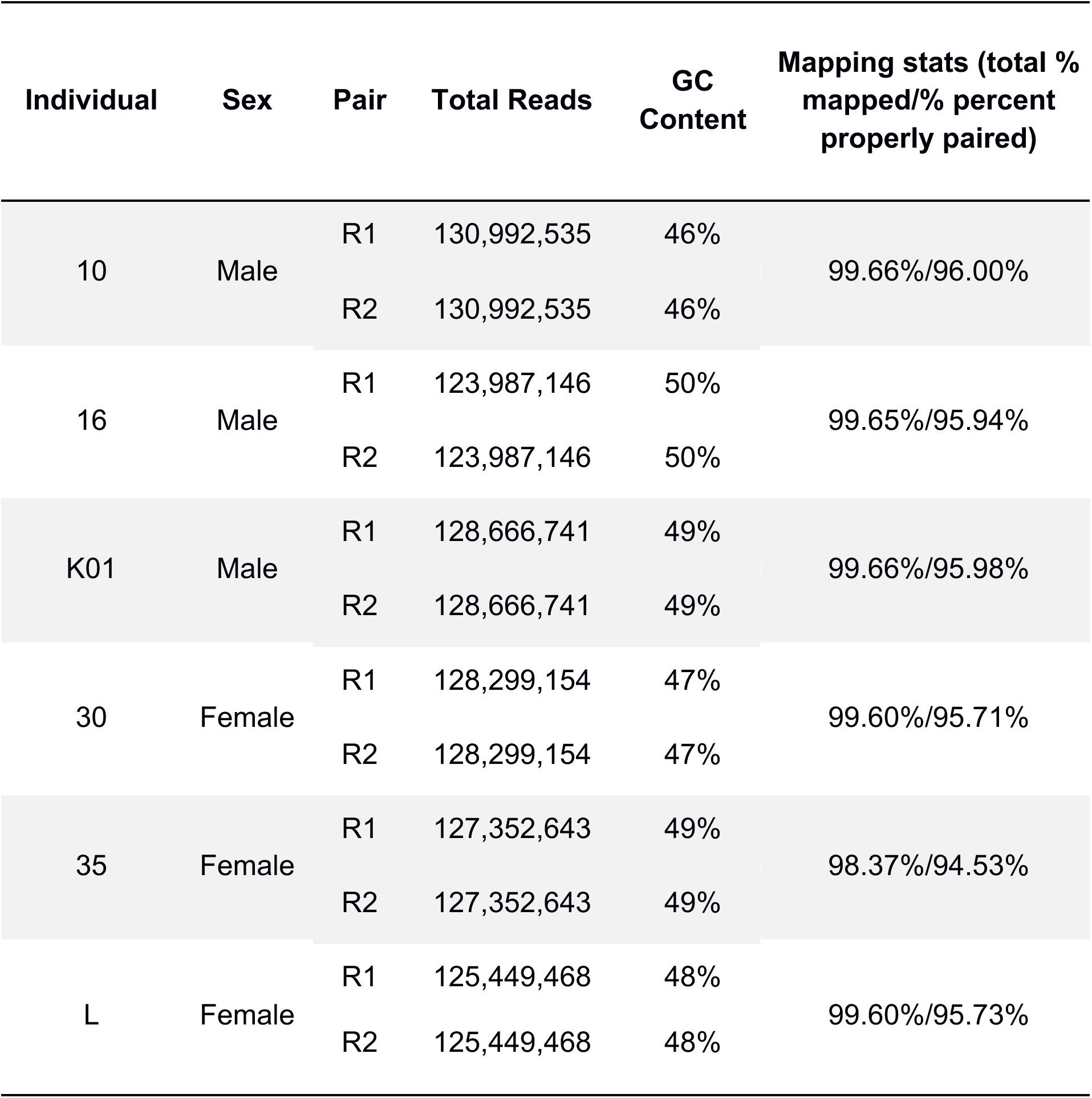
DNA Resequencing Sequencing Statistics for Gila Monster. Read stats calculated using FastQC, mapping stats calculated using sambamba flagstat.

| Individual | Sex | Pair | Total Reads | GC Content | Mapping stats (total % mapped/% percent properly paired) |
| --- | --- | --- | --- | --- | --- |
| 10 | Male | R1 | 130,992,535 | 46% | 99.66%/96.00% |
|  |  | R2 | 130,992,535 | 46% |  |
| 16 | Male | R1 | 123,987,146 | 50% | 99.65%/95.94% |
|  |  | R2 | 123,987,146 | 50% |  |
| K01 | Male | R1 | 128,666,741 | 49% | 99.66%/95.98% |
|  |  | R2 | 128,666,741 | 49% |  |
| 30 | Female | R1 | 128,299,154 | 47% | 99.60%/95.71% |
|  |  | R2 | 128,299,154 | 47% |  |
| 35 | Female | R1 | 127,352,643 | 49% | 98.37%/94.53% |
|  |  | R2 | 127,352,643 | 49% |  |
| L | Female | R1 | 125,449,468 | 48% | 99.60%/95.73% |
|  |  | R2 | 125,449,468 | 48% |  |

**Supplemental Table S2:** RNA Sequencing Statistics for Gila Monster. Read stats calculated using FastQC, mapping stats calculated using sambamba.

| Individual | Sex | Pair | Total Reads | GC Content | Mapping stats (total % mapped/% percent properly paired) |
| --- | --- | --- | --- | --- | --- |
| 10 | Male | R1 | 75,069,664 | 46% | 94.21%/91.54% |
|  |  | R2 | 75,069,664 | 46% |  |
| 16 | Male | R1 | 66,659,537 | 50% | 84.35%/80.59% |
|  |  | R2 | 66,659,537 | 50% |  |
| K01 | Male | R1 | 65,347,060 | 49% | 87.04%/82.70% |
|  |  | R2 | 65,347,060 | 49% |  |
| 30 | Female | R1 | 61,087,539 | 47% | 91.51%/88.05% |
|  |  | R2 | 61,087,539 | 47% |  |
| 35 | Female | R1 | 72,499,762 | 49% | 83.95%/79.45% |
|  |  | R2 | 72,499,762 | 49% |  |
| L | Female | R1 | 70,669,648 | 48% | 90.28%/86.36% |
|  |  | R2 | 70,669,648 | 48% |  |

**Supplemental Table S3.**
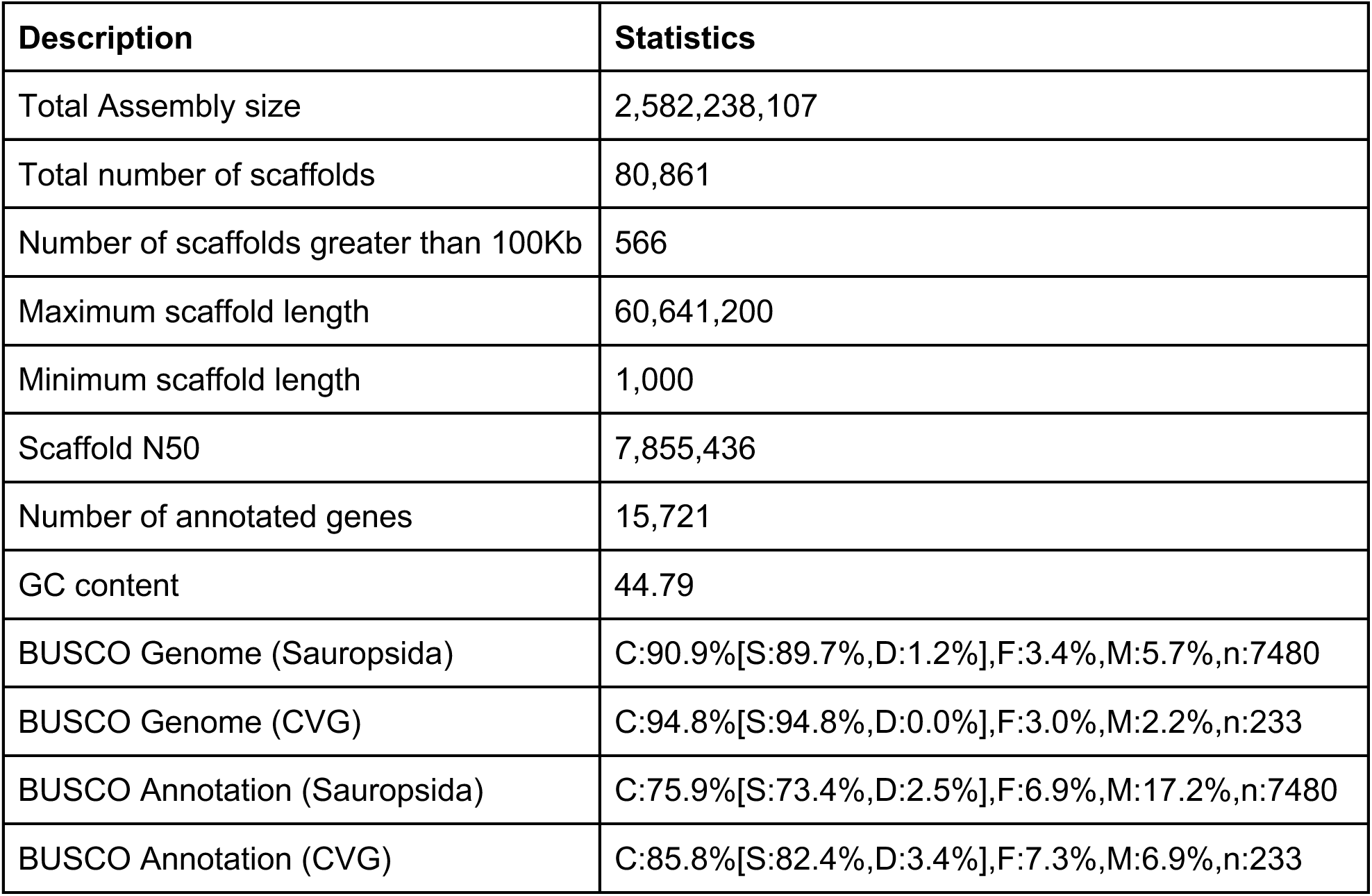
Gila monster (*Heloderma suspectum,* male #10) Genome Statistics, part 1.

**Supplemental Table S4.** Gila monster (*Heloderma suspectum*, male #10) genome statistics, part 2.

| | Scaffolds ≥ 1bp | Scaffolds ≥ 10kb | Scaffolds ≥ 50kb |
| --- | --- | --- | --- |
| Total length | 2.58 Gb | 2.31 Gb | 2.13 Gb |
| Number of scaffolds | 80,861 | 12,703 | 913 |
| Number of contigs | 152,240 | 73,769 | 57,630 |
| Scaffold N50 | 7.855 Mb | 9.23 Mb | 9.93 Mb |
| Scaffold L50 | 94 | 78 | 68 |
| Contig N50 | 35.49 Kb | 42.6 Kb | 47.8 Kb |
| Contig L50 | 15,710 | 12,233 | 10,298 |
| Max scaffold length | 60.64 Mb | 60.64 Mb | 60.64 Mb |
| Max contig length | 469.1 Kb | 469.1 Kb | 469.1 Kb |

**Supplemental Table S7.**
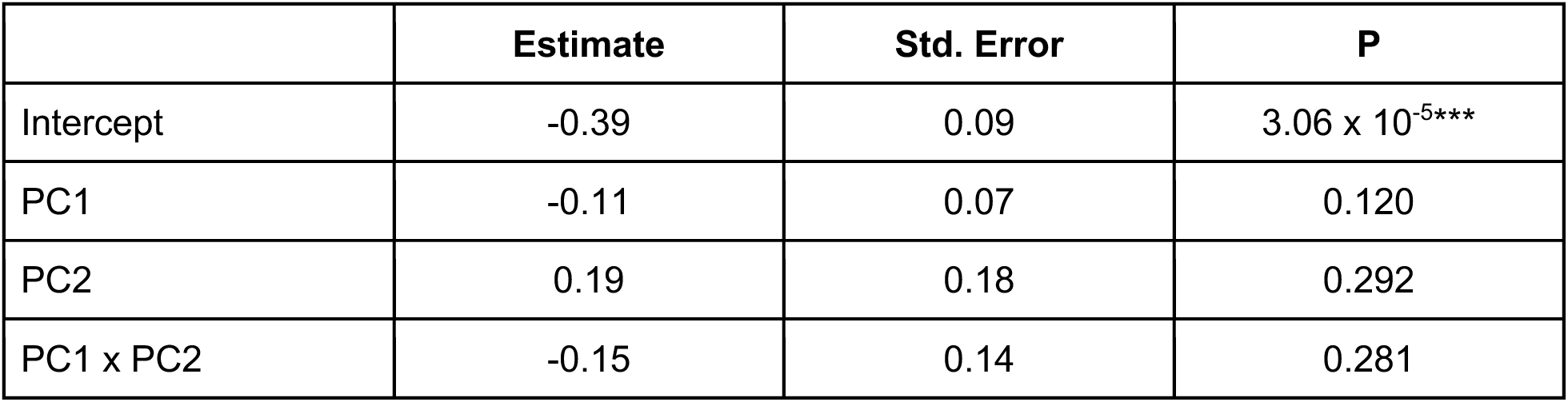
Results of linear model testing the effect of selection intensity on dosage balance. PC1 corresponds to sexually concordant selection (larger values indicate more concordance) and PC2 captures sex-biases in selection intensity (larger values indicate a male bias and smaller values indicate a female bias).

**Supplemental Figure S1:**
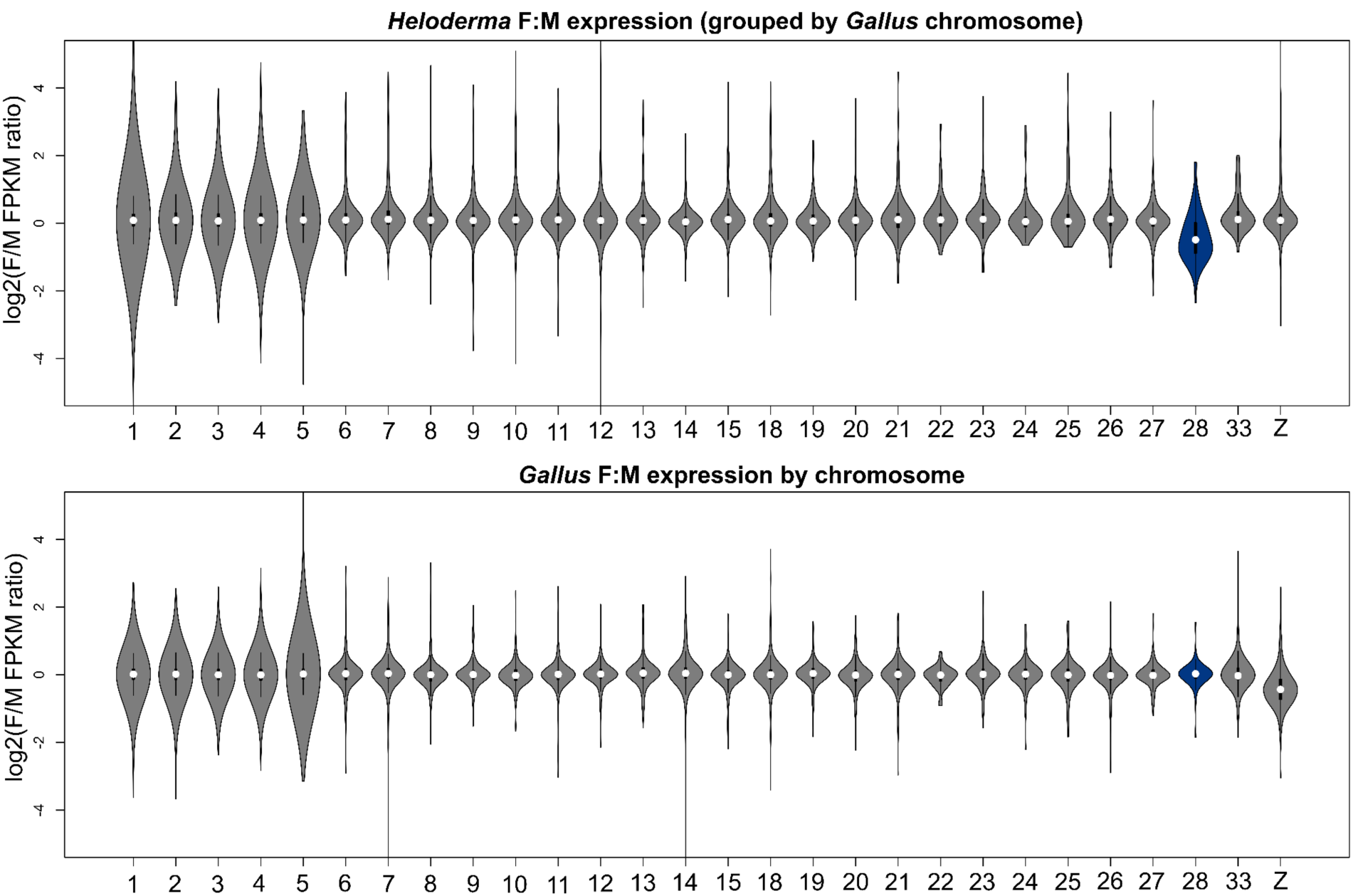
F/M gene expression grouped by chicken chromosomes. Log2(F/M FPKM ratios) for genes clustered by their orthologous position in *Gallus* for *Heloderma suspectum* (top) and *Gallus gallus* (bottom), highlighting the drop in F/M expression in gene orthologous to Gg28 in *Heloderma*.

**Supplemental Figure S2:**
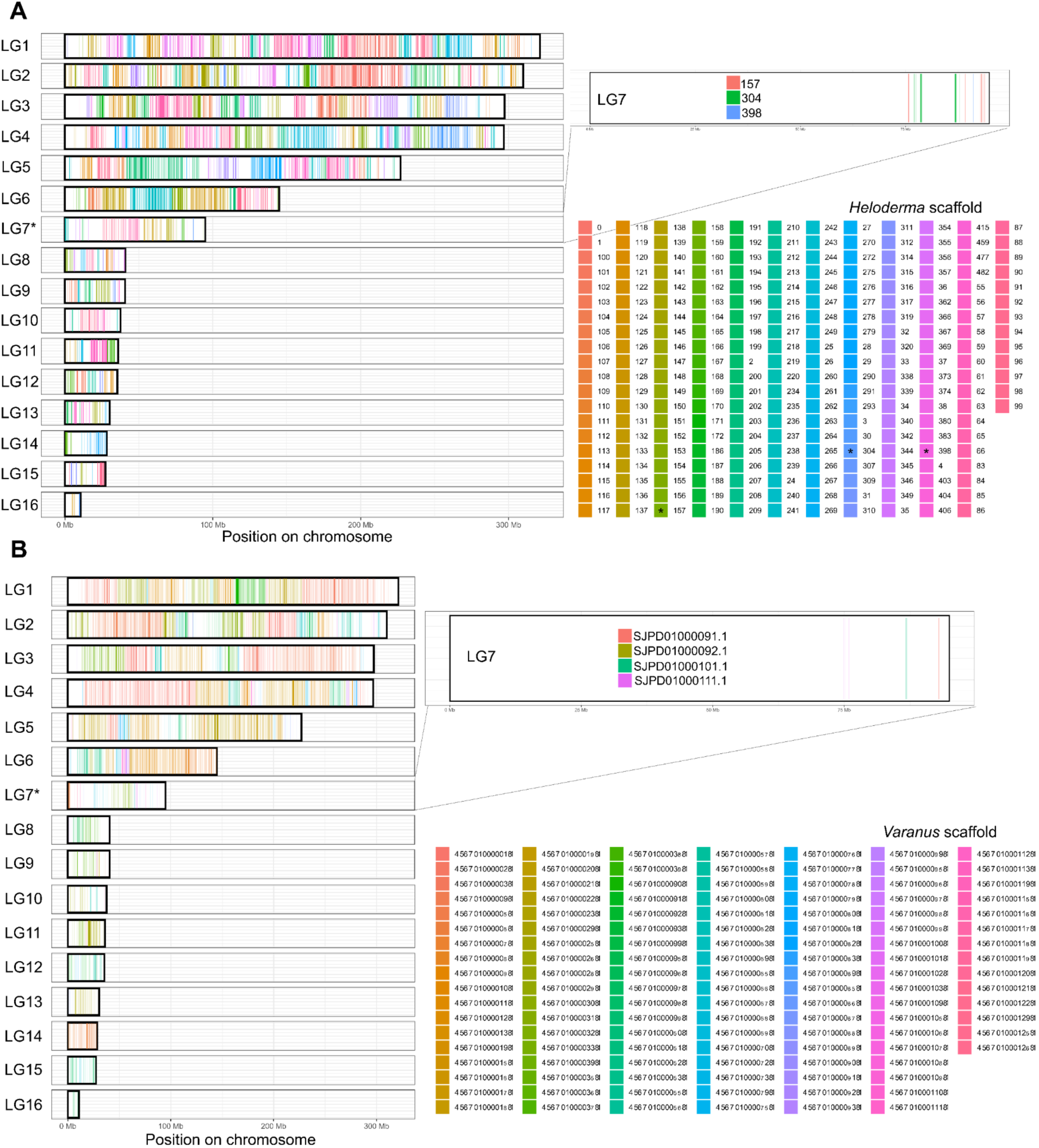
*Heloderma suspectum* and *Varanus komodoensis* genomes aligned to the *Shinisaurus crocodilurus* genome. (A) Sex-linked scaffolds in *H. suspectum* mapping to the distal region of *S. crocodilurus* chromosome 7 (LG7). (B) Sex-linked scaffolds in *V. komodoensis* also mapping to the distal region of *S. crocodilurus* LG7.

**Supplemental Figure S3:**
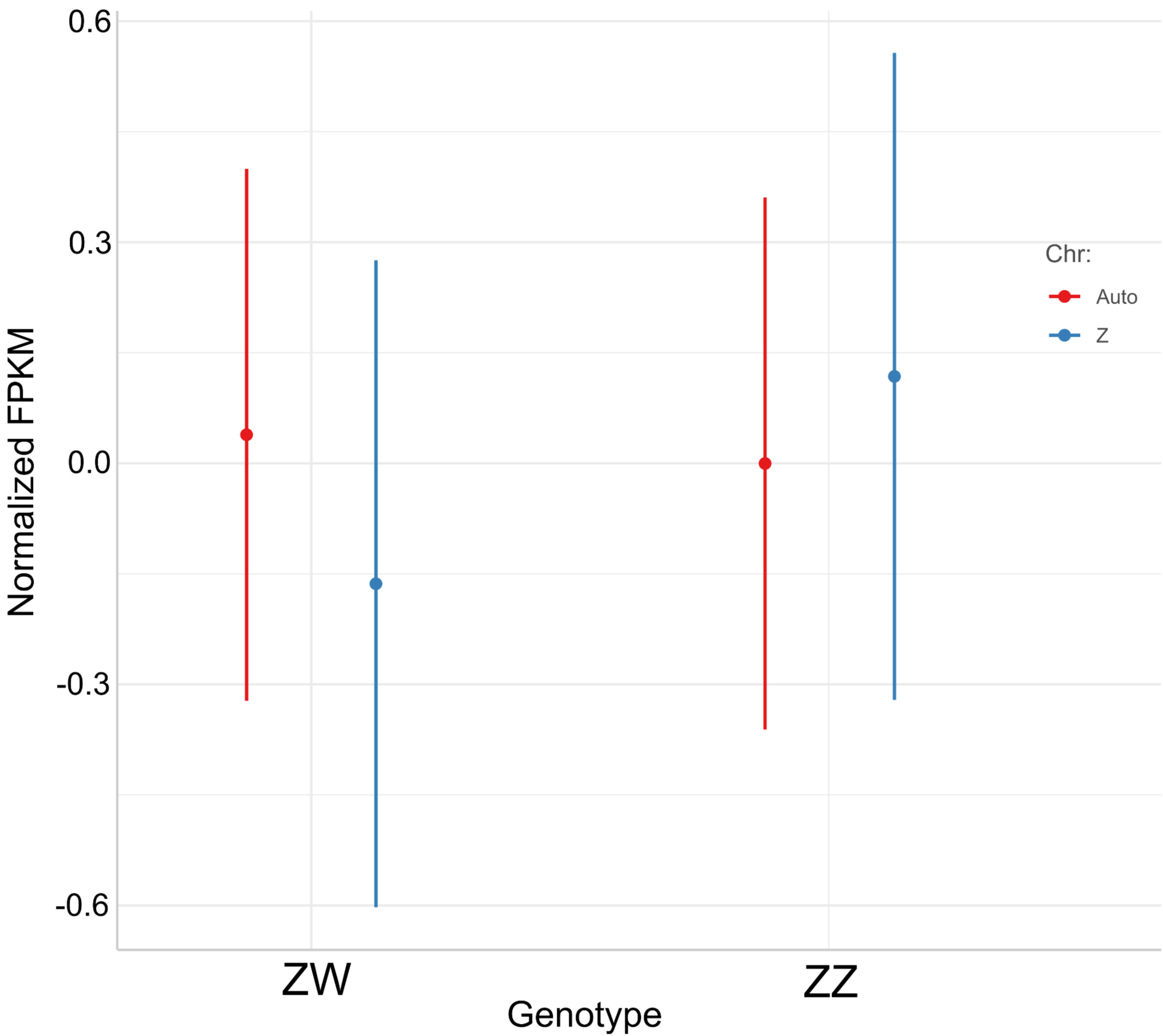
Marginal effects of the interaction between sex (ZZ: male; ZW: female) and chromosome type (autosome vs. Z chromosome) on expression in Gila monster. Estimates from the full model for dosage compensation using chicken as an outgroup. Fixed effects were sex, Z-linkage, and their interaction, while transcript ID, individual ID, and the interaction between male and female expression in chicken were included as random effects. Expression in chicken serves as a proxy for expression in the ancestral autosomal condition.

**Supplemental Figure S4.**
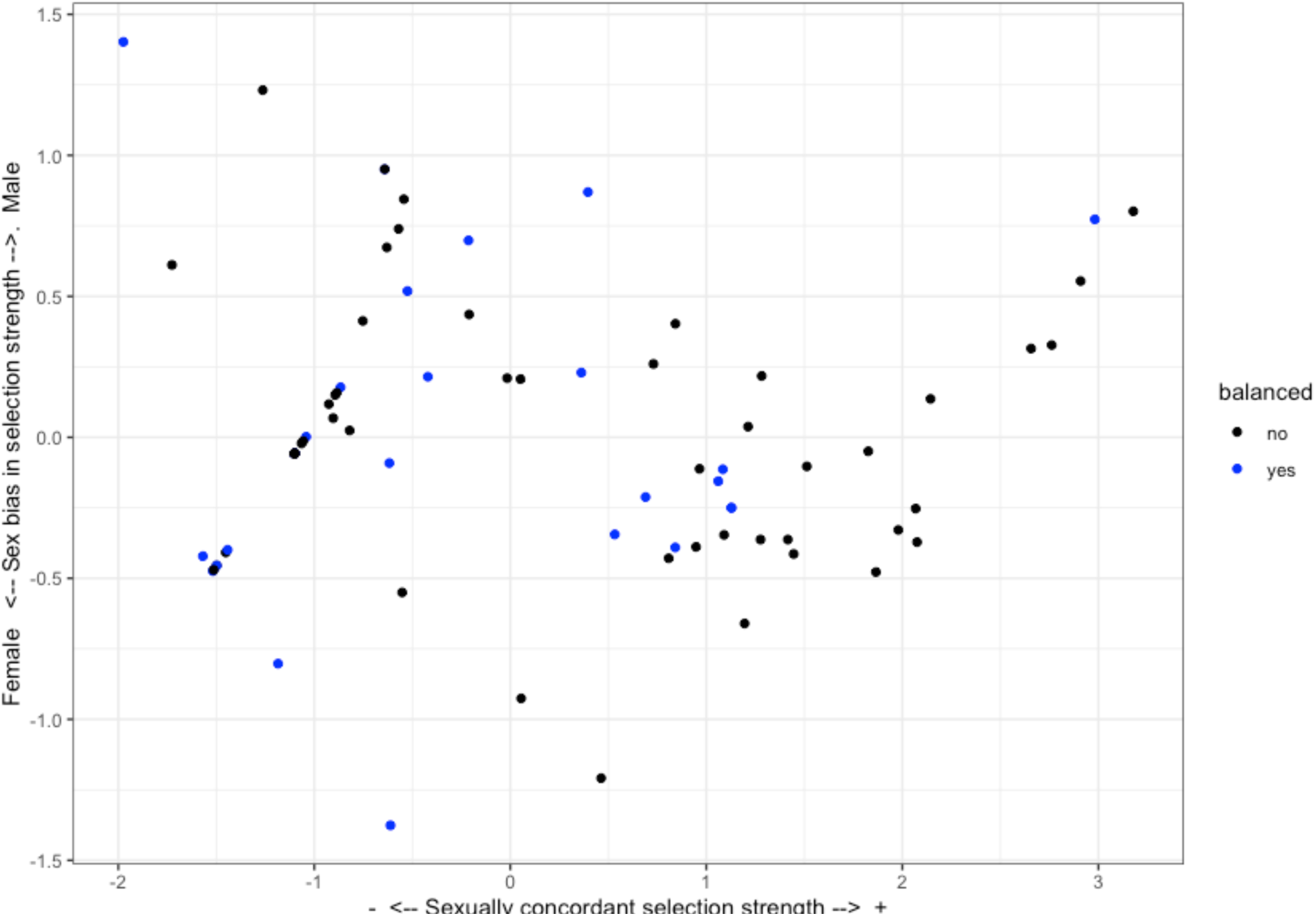
Selection and dosage balance in Gila monster. Illustrating the effects of sexually concordant selection (x-axis) and sex-biased expression (y-axis) on dosage balance. For this figure, transcripts are marked as balanced or female-biased (blue) if the log2 ratio of female to male expression is greater than –0.32 (equivalent to a raw ratio of approximately 0.8 or greater). On the x-axis, larger values indicate stronger sexually concordant selection. On the y-axis, more positive values are associated with a greater male bias, and more negative values are associated with a female bias.

